# Archaeogenomics of a ~2,100-year-old Egyptian leaf provides a new timestamp on date palm domestication

**DOI:** 10.1101/2020.11.26.400408

**Authors:** Oscar A. Pérez-Escobar, Sidonie Bellot, Muriel Gros-Balthazard, Jonathan M. Flowers, Mark Nesbitt, Philippa Ryan, Rafal M. Gutaker, Tom Wells, Rowan Schley, Diego Bogarín, Natalia Przelomska, Steven Dodsworth, Rudy Diaz, Manuela Lehmann, Peter Petoe, Wolf L. Eiserhardt, Michaela Preick, Michael Hofreiter, Irka Hajdas, Alexandre Antonelli, Ilia J. Leitch, Barbara Gravendeel, Maria Fernanda Torres, Guillaume Chomicki, Susanne S. Renner, Alexander S.T. Papadopulos, Michael Purugganan, William J. Baker

**Author notes:** **Correspondence** (OAPE) or (WJB).

## Abstract

- The date palm (*Phoenix dactylifera*) has been a cornerstone of Middle Eastern and North African agriculture for millennia. It is presumed that date palms were first domesticated in the Persian Gulf and subsequently introduced into North Africa, where their evolution in the latter region appears to have been influenced by gene flow from the wild relative *P. theophrasti*, which is restricted to Crete and Turkey. However, the timing of gene flow from *P. theophrasti* to *P. dactylifera* remains unknown due to the limited archaeobotanical evidence of *P. theophrasti* and their exclusion from population genomic studies.
- We addressed this issue by investigating the relatedness and ancestry of a ~2,100-year-old *P. dactylifera* leaf from Saqqara (Egypt), combining genome sequencing of this ancient specimen with a broad sample of date palm cultivars and closely related species.
- The ancient Saqqara date palm shares close genetic ancestry with North African date palm populations. We find clear genomic admixture between the Saqqara date palm, *P. theophrasti* and the closest known relative *P. sylvestris*.
- Our study highlights that gene flow from *P. theophrasti* and *P. sylvestris* to North African date palms had already occurred at least ~2,100 years ago, providing a minimum timestamp for hybridisation between species.

## Introduction

The start of plant crop domestication some 10,000-12,000 years ago was arguably one of the most important events in human history (Diamond, 2002). The domestication of plant crop and husbandry of animals allowed the sustained nutrition of large sedentary human population settlements (Fuller et al., 2014; Larson et al., 2014; Richter et al., 2017; Arranz-Otaegui et al., 2018). Elucidating the domestication history of major plant crops is thus an important scientific challenge, which requires collaboration between scholars of archaeology, anthropology, taxonomy, genetics and genomics. The widespread availability of high-throughput DNA sequencing has revolutionized the study of plant crop domestication history, leading to many unpreceded insights, such as the identification of crop progenitors (Ling et al., 2013; Gros-Balthazard et al., 2017, Chomicki et al. 2020), hybridization and introgression events following the origin of crops (e.g. Cornille et al., 2012; Hufford et al., 2012; Baute et al., 2015; Muñoz-Rodrígez et al., 2018), refinement of the geographic origins of crops (Besnard et al., 2017; Cubry et al., 2018), the identification of genes controlling key domestication traits (Zhou et al., 2016; Stitzer and Ross-Ibarra, 2018) and more generally of convergent evolutionary processes that have challenged orthodoxies on domestication (reviewed by Purugganan, 2019). The application of genomic approaches to crop wild relatives is also bringing critical new resources for crop improvement (reviewed by Brozynska et al., 2016).

With more than 8 million tonnes of fruits produced yearly, the date palm (*Phoenix dactylifera* L.) has been a cornerstone of Middle Eastern and North African agriculture for millennia. Despite its economic importance, the date palm domestication history is far from well understood (reviewed in Gros-Balthazard et al., 2018). Archaeological evidence, ancient texts and iconographies all point to the use of date palms for millennia in North Africa, the Middle East and as far as Pakistan (Tengberg, 2012). The first evidences of cultivation date to the end of the 4^th^ millennium B.C.E. in the Persian Gulf region (reviewed in Tengberg, 2012). It is thus presumed that date palms were first domesticated in this region and subsequently introduced into North Africa (reviewed by Gros-Balthazard et al., 2018). However, a study that resequenced 62 date palm cultivars from the Middle East and North Africa found more genetic diversity in North African date palm populations, challenging the simple Middle-Eastern origin hypothesis (Hazzouri et al., 2015). Population genomic analyses of date palm cultivars and other *Phoenix* species revealed extensive introgressive hybridization of the North African date palm with *P. theophrasti* Greuter - up to 18% of the genome of North African cultivars was shared with the Cretan date palm (Flowers et al., 2019). Clearly, date palm evolution in North Africa has been influenced by gene flow from the wild relative *P. theophrasti*. However, the timing of this introgressive event is unknown.

To address this important knowledge gap in date palm domestication, we sequenced the genome of a ~2,100-year-old date palm (*P. dactylifera* L.) leaf found in Saqqara, Egypt, radiocarbon-dated to the Late Period of ancient Egypt (357-118 B.C.E.). We found that the genomic ancestry of the ancient Saqqara date palm can be traced to modern domesticated North African *P. dactylifera* and the close wild relatives *P. sylvestris* and *P. theophrasti*. Hybridisation and gene flow between North African *P. dactylifera, P. theophrasti* and *P. sylvestris* had already taken place by ~2,100 years ago. Our study thus provides a minimum bound on the timing for gene flow between date palms and their close living relatives.

## Materials and Methods

### Plant taxon sampling

Our sampling builds upon recent population genomic studies of date palms by Flowers et al. (2019) and Gros-Balthazard (2017) with a total of 36 individuals representing seven species. We sampled seventeen individuals of wild and cultivated Asian and North African *P. dactylifera* populations, as well as 18 accessions of five closely related species, namely *P. atlantica*, *P. canariensis, P. reclinata, P. sylvestris* (the sister species of the date palm) and *P. theophrasti* following the current accepted taxonomy of the genus *Phoenix* (Barrow, 1998; Gros-Balthazard et al., 2020a). In addition, a shallow genomic representation of the New Guinean palm *Licuala montana* was sequenced to produce a plastid genome assembly that was subsequently used as the root for phylogenetic analyses. This species was chosen due to the sister relationship between the palm tribes Phoeniceae (containing *Phoenix*) and Trachycarpeae (containing *Licuala*) (Baker and Dransfield, 2016).

Whole genome sequence data from these taxa was obtained from the NCBI sequence read archive repository. Twenty million reads were downloaded for each accession using the tool *fastq-dump* of the SRA toolkit. Nearly all accessions sampled are linked to vouchers and have known origins (Gros-Balthazard et al., 2017; Flowers et al., 2019); detailed information on their provenance, average read length and number of bases downloaded are provided in Tables S1 and S2.

### Radiocarbon dating and ancient DNA extraction

To determine with accuracy the age of the Saqqara date palm item (accession EBC 26796), one cm^2^ of leaf removed from the edge of the sample was sent to the Laboratory of Ion Beam Physics, ETH-Zurich. The leaf sample underwent a treatment with solvents and an acid–base–acid washes (Hadjas et al., 2008) to remove potential contamination of waxes, carbonates and humic acids. The dry, clean material (weighing 2.6 mg, equivalent to 1 mg of carbon) was weighed into tin cups for combustion in the Elemental Analyser for subsequent graphitization (Němec et al., 2010). The resulting graphite was pressed into aluminium cathodes and the ^14^C/^12^C and ^13^C/^12^C ratios were measured using the Mini Carbon Dating System dedicated accelerator mass spectrometry facility (Synal et al., 2007). The radiocarbon age was calculated following the method described by Stuiver et al. (1977) using the measured ^14^C content after correction for standards, blank values and fractionation (δ13C values were measured semi-simultaneously on graphite). The reported conventional age in years BP (before 1950 AD or CE) was calibrated to a calendar age using OxCal version 4.2.4 (Reimer et al., 2013) and the IntCal13 atmospheric curve (Bronk-Ramsey, 2013).

Ancient DNA (aDNA) was extracted by grinding a small piece of a leaflet (<1 cm^2^, 5.8 mg) with a Retsch mill (MM 400). DNA extraction was performed following the modified protocol of Wales et al. (2014) (Pedersen et al., 2014; Dabney et al., 2013). For the digestion treatment, a lysation buffer containing 0.5 % (w/v) N-lauroylsarcosine (Sigma Aldrich L9150-50G), 50 mM Tris-HCl (Thermo Fisher Scientific 15568025), 20 mM EDTA (VWR E177-500MLDB) 150 mM NaCl (Thermo Fisher Scientific AM9760G), 3.3 % 2-mercaptoethanol (Sigma Aldrich 63689-25ML-F), 50 mM DL-dithiothreitol (Sigma Aldrich D9779-250MG) and 0.25 mg/mL Proteinase K (Promega V3021) was applied to the leaflet powder as described in Wales et al. (2014). DNA purification was performed according to Dabney et al. (2013) but with reduced centrifugation speed (450 x *g*), following Basler et al. (2017).

### Ancient DNA library preparation and sequencing

A genomic Illumina library was prepared from the extracted aDNA following the single-stranded protocol of Korlević et al. (2015). The protocol included the treatment with Uracil-DNA-Glycolase (New England Biolabs M0279) to remove uracil residues and Endonuclease VIII (New England Biolabs M0299) to cleave DNA strands at abasic sites. Circligase II (2.5 U/μl; Biozym 131406) was used for the fill-in reaction which was carried out overnight. A quantitative PCR was performed on a PikoReal 96 Real-Time PCR machine (Thermo Fisher Scientific TCR0096) using 0.2 % of the unamplified library and the following thermal profile: 10 min initial denaturation step at 95 °C, followed by 40 cycles of: 15 s at 95 °C, 30 s at 60 °C, and 1 min at 72 °C. The quantitative PCR reaction mix contained a final volume of 10 μL: 1 μL of diluted library, 1 x SYBR Green qPCR Master Mix (Applied Biosystems 4309155), 0.5 μM of each primer IS7 and IS8. Three replicates of each library were used. Indexing PCR was performed by the appropriate number of cycles according to the results of the qPCR, with 8 bp indices added to the 5’ and 3’ adapters. The PCR and final concentrations used were the same as described by Gansauge and Meyer (Korlević et al., 2017), but with a final volume of 80 μL using 20 μL of template. DNA sequencing was performed on an Illumina NextSeq 500 sequencing platform, using the 500/550 High Output v2 kit (75 cycles, Illumina FC-404-2005), with a custom read-1 (Perdersen et al., 2014) and a custom index-2 (Paijmans et al., 2017) sequencing primer. All extractions and library preparations were performed in the ancient DNA facility of the University in Potsdam; negative controls were included in all steps.

### Genome skimming of *Licuala montana*

We extracted genomic DNA from silica-dried leaf tissue of *L. montana* using the Qiagen DNeasy Plant kit, following the manufacturer’s protocol. A genomic Illumina paired-end library was prepared using the NEBNext Ultra II library preparation kit, following the manufacture’s protocol and with an average insert size of 150 bp. Library sequencing was performed by the company Genewiz (New Jersey, USA) on a HiSeq platform. A total of 4 million paired-end reads was produced.

### High-throughput read data processing

The Illumina raw reads were quality filtered using Trim Galore v.0.4 (Krueger, 2015), discarding sequences with an averaged phred33 score below 20. Pre- and post-trimming read quality was assessed using FASTQC v.0.1 (Andrews et al., 2015). The proportion of endogenous DNA sequence present in the ancient Saqqara date palm leaf extract was assessed by blasting the trimmed read data against the nuclear and plastid genomes of *Phoenix dactylifera* (Khalas variety, assembly GCA000413155.1 [Al-Mssallem et al., 2013]) using BLAST+ v.2.8.1 (McGinnis and Madden, 2013), an e-value of 0.001 and a coverage threshold of 80%. We then determined the proportion of nuclear/plastid ancient date palm read data sequenced by filtering the number of hits mapped by blast onto nuclear, plastid and mitochondrial scaffolds.

Given the low proportion of nuclear genomic data recovered from the ancient Saqqara date palm leaf (see *Results*), we mapped trimmed reads of both modern and ancient accessions on nuclear and plastid targeted scaffolds (i.e. scaffolds with Saqqara date palm leaf reads mapped). These represented 198 contigs, or 26.8% (149.01 Mb) of the *P. dactylifera*’ nuclear genome assembly. We investigated the proportion and position of mis-incorporated nucleotides in the Saqqara date palm leaf DNA using aligned aDNA reads and the tool mapDamage2 v.2.0.9 (Jónsson et al., 2013). We compared nucleotide mis-incorporation patterns between the aligned aDNA reads of the Saqqara date palm leaf and DNA reads of modern date palm accessions (SRR5120110). Finally, to reduce the fraction of mis-incorporated nucleotides mapped onto the reference genome, we trimmed two bases at the 3’ and 5’ end of the aDNA reads, using Trim Galore v.0.4. Read mapping, alignment and DNA damage analyses were implemented through the pipeline PALEOMIX v.1.2.13 (Schubert et al., 2014). The trimmed read data were mapped using the software bowtie v.2.3.4.1, followed by a realigning step around indels and filtering of duplicated reads with the software GATK v.3.8.1 (McKenna et al., 2010) and Picard-tools v.1.137 (Thomer et al., 2016). Mis-incorporated nucleotides in DNA fragments are characteristic of sequence data derived from historical and archaeobotanical specimens (Estrada et al., 2018). Read mapping, and average coverage statistics for each accession sampled in this study are provided in Table S1.

To account for biases in the mapping of aDNA read data onto the reference genome (Günther and Nettelblad, 2019) and test the robustness of our population genomic inferences against missing data (Skotte et al., 2013), we also mapped the ancient and modern DNA reads onto 18 highly contiguous scaffolds of a newly assembled nuclear genome of *P. dactylifera* (four-generations backcross of a Bahree cultivar, assembly CA0009389715.1 [Hazzouri et al., 2019]), representing 50% of the nuclear genome (~380 Mb). Read mapping and alignment were conducted using the same procedure and tools as specified above.

### Plastid phylogenomic analyses of *Phoenix*

We produced consensus plastid genome sequences of modern date palm accessions from the BAM files produced by PALEOMIX by following a modified statistical base-calling approach of Li et al. (2008), i.e. minimum depth coverage of 10, and bases matching at least 50% of the reference sequence. Because the attained average coverage of the Saqqara date palm leaf plastid genome was ~2x (Table S1), the consensus plastid sequence for this accession was produced by using a minimum depth coverage of 2, bases matching at least 50% of the reference sequence and missing data represented as Ns whenever parts of the reference plastid genome were not covered by aDNA reads. The whole plastid genome consensus sequences were produced in Geneious v.8.0. Consensus plastid genome sequences were aligned with Mauve using a progressive algorithm and assuming collinearity (Darling et al., 2004). The resulting ~150,000 bp alignment was first trimmed to exclude mis-aligned regions and positions with >90% missing data (final alignment length of 103,807 bp) and then subjected to Maximum Likelihood (ML) tree inference in RAxML v8.0 (Stamatakis, 2014), using the GTR substitution model, 25 gamma rate categories, and 1,000 bootstrap replicates.

### Population structure and nuclear phylogenetic position of the Saqqara date palm

Given low-depth high-throughput sequencing data produced for the ancient Saqqara date palm leaf, we relied on genotype likelihoods (GL) to place the aDNA nuclear genomic data of the Saqqara date in range with genomic sequences of modern *Phoenix* samples. We computed nuclear GL using the software ANGSD v.0.929 (Korneliussen et al., 2014), by implementing the GATK GL model, inferring the minor and major alleles, and retaining polymorphic sites with a minimum p-value of 1-e^6^. To reveal the relationship of the ancient Saqqara date leaf nuclear genome to the genetic diversity of modern samples of *Phoenix*, we conducted principal component (PCA) and population structure analyses using nuclear GLs and the tools PCangds (Meisner and Albrechtsen, 2018), and NGSadmix (Skotte et al., 2013) of the software ANGDS, respectively. Because PCA can be particularly affected by the proportion of overlapping sites between modern and ancient populations (Ausmees, 2019), we computed covariance matrices by using only GLs derived from sites that were shared across all the modern individuals and the ancient Saqqara date palm leave (i.e. option -minInd set to 35), and a maximum of 1000 iterations. Admixture analyses were conducted with number of population (K) set from two to eight and a maximum of 20,000 iterations. The best K was selected by comparing the resulting likelihood values derived from each K iteration.

To test for admixture between the Saqqara date palm and other lineages amongst date palms or closely related species (*P. atlantica*, *P. sylvestris* and *P. theophrasti*), we used the D-statistic framework as implemented in the software ANGSD. To account for the differences in sequencing coverages obtained for modern individuals and the Saqqara leaf and to assess the robustness of our introgression tests, two different approaches were followed, namely a) between nuclear genomes of each individual (i.e. sampling one base from reads of one individual per population [Korneliussen et al., 2014]) and b) between populations (i.e. considering all reads from multiple individuals in each population [Soraggi et al., 2018]). Nuclear GL were used as input and D-statistics for both tests. In (a) D-statistics were calculated by sampling a random base at each analysed position in blocks of 5 million bp, removing all transitions to rule out possible post-mortem base misincorporations in the Saqqara sample together with reads with qualities lower than 30, and setting *P. reclinata* as a fixed outgroup terminal following the same experimental design as in Flowers et al. (2019). In (b) individuals were assigned to six populations defined according to their species identity, e.g. all modern individuals of *P. dactylifera* were assigned to one population (Table S2); the Saqqara leaf was assigned to its own population as well as the outgroup (*P. reclinata*), following the recommendations provided by Soraggi et al. (2018). D-statistics were then calculated by sampling reads from multiple individuals in each population. Moreover, to account for the influence of polymorphisms in the outgroup taxon, we executed population tests using polymorphic and non-polymorphic sites (Table S5) and only non-polymorphic sites in the outgroup (i.e. “–enhance” flag in ANGSD; Table S6). The significance of the analyses was assessed by executing a block-Jacknife test which derived standard errors and Z–scores. For population level analyses, *p-values* were derived. The admixture of the Saqqara date palm was discussed based solely on significant *p*- and D-statistic values (i.e. |Z| > 3). D-statistic values, standard deviations, Z-scores and the number of evaluated sites for each topological permutation are provided in Tables S3A-C. NGSadmix and D-statistic analyses were conducted on filtered GLs derived from read data mapped on: a) 198 contigs representing 26.8% of the *P. dactylifera* genome (assembly GCA000413155.1); and b) 18 scaffolds representing 50% of the nuclear genome of *P. dactylifera* (assembly GCA0009389715.1, see *High-throughput read data processing* section of *Methods* above).

## Results

### DNA sequencing of an archaeological date palm leaf from Saqqara

Our archaeological sample is from an object made from date palm leaflets discovered in the temple complex of the animal necropolis of Saqqara, an Egyptian UNESCO World Heritage site located 20 km south of Cairo and adjacent to the Nile valley. The object, currently held in the Economic Botany Collection at the Royal Botanic Gardens, Kew, was recovered during the 1971-2 excavation season from the ‘West Dump’. The site is a mixed refuse deposit dating between 500-300 B.C.E. that also contained other objects such as possible ‘brushes’ made from date palm, papyri, jar-stoppers, amulets and other debris including seeds (Martin et al., 1981). The object consists of a plaited portion of a leaf (including leaflets and rachis) and was originally considered as a ‘head-pad’ by excavators, however there are no analogous finds from other sites supporting this interpretation. A virtually identical object from the ‘West Dump’ at Saqqara is held in the British Museum collections (accession EA68161) where it is identified as possibly ‘part of the lid of a basket’. We speculate that the object, hereafter referred to as “Saqqara leaf” (Fig. 1), may instead have been part of the layering used to close and seal a vessel, similar to those found in the New Kingdom (1570–1070 B.C.E) (Hope, 1977). We radiocarbon-dated the Saqqara leaf to 2,165 ± 23 BP (ETH-101122), or to a calibrated date of 357-118 B.C.E, thus confirming its burial during the Late Period or the Ptolemaic Kingdom of ancient Egypt.

**Figure 1.**
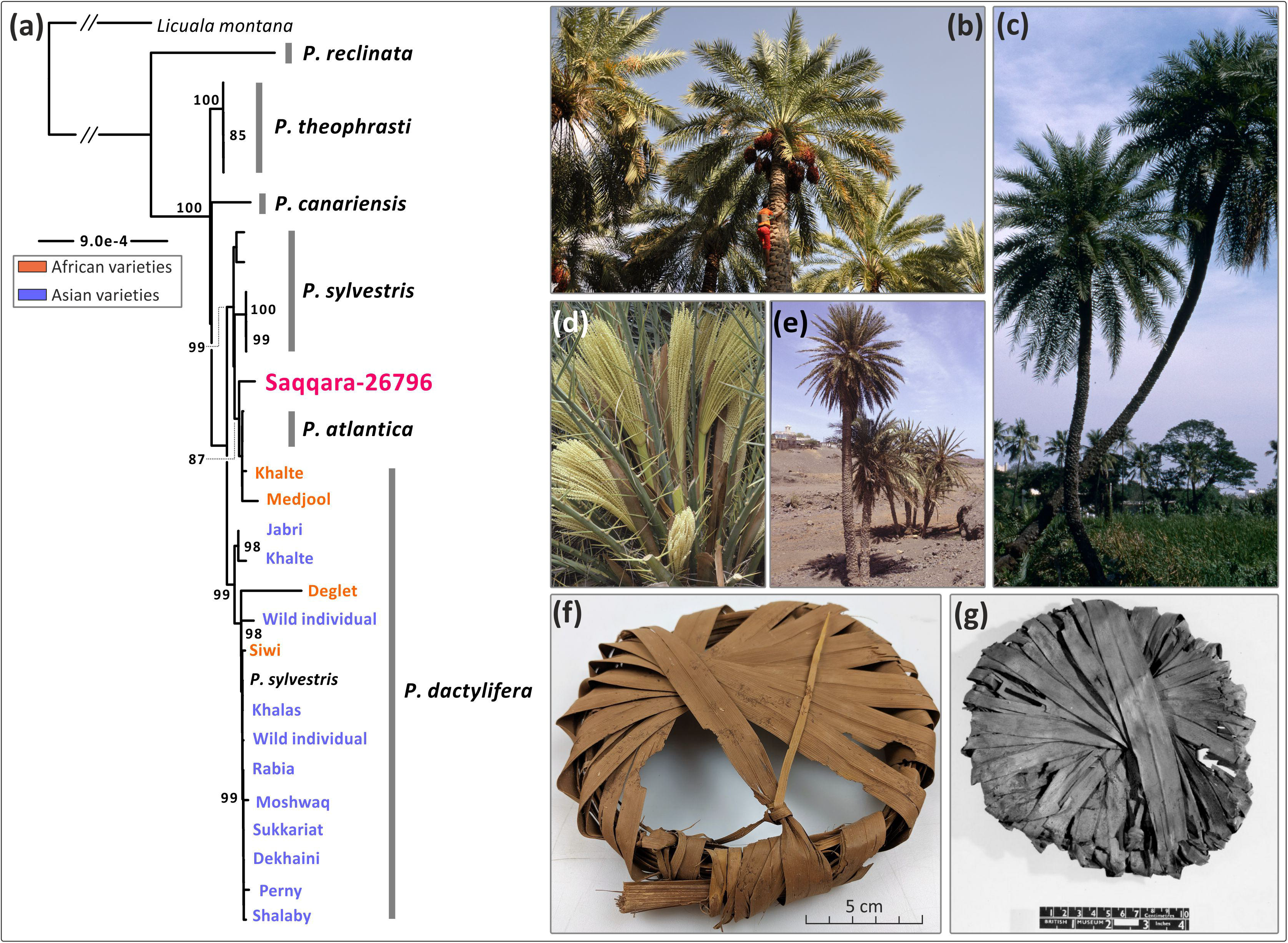
Phylogenetic placement of the Saqqara specimen amongst *Phoenix* species. (**a**) Maximum Likelihood analysis of whole plastome sequences showing the placement of the Saqqara specimen amongst *Phoenix* species (accession numbers for each terminal are provided in Fig. S2). (**b**) Individual of *Phoenix dactylifera* bearing fruits. (**c**) Individual of the sugar date tree (*P. sylvestris*), the closest known leaving relative of *P. dactylifera*. (**d**) Male inflorescence of *P. theophrasti*. (**e**) Individuals of *P. atlantica*. (**f**) Saqqara 26796, a jar-stopper made of date palm leaflets (excavation inventory number 102, Kew Economic Botany Collection number 26796). (**g**) A similar object to the Saqqara specimen number 26796, also made of date palm leaflets thought to be a basket-lid and found in Saqqara. Photos: Penelope Dawson (**b**), Sasha Barrow (**c**), John Dransfield (**d**), William J. Baker (**e**), Mark Nesbitt (**f**), © The Trustees of the British Museum CC BY-NC-SA 4.0 (**g**).

We sequenced ~400 million reads from the Saqqara leaf of which up to ~4% (i.e. *c.* 16 million reads) were identified to be from endogenous DNA of *P. dactylifera* (Table S1). As expected from ssDNA libraries, nucleotide misincorporations (C to T), which are indicative of DNA damage, predominantly occurred towards both ends of the reads, visible even after an uracil reduction procedure. Read length distributions were centred on 35 bp (Fig. S1), consistent with sequencing data from similarly-aged material (Scott et al., 2019; Ramos-Madrigal et al., 2016). Although the average read depth was only ~2x, we obtained a near-complete representation of the plastid genome of the Saqqara sample, covering 95% of the plastome of modern date palms (see *Methods*). We also recovered up to ~755 million base pairs of the *P. dactylifera* nuclear genome (Table S1).

### Phylogenetic placement of the Saqqara leaf and detection of introgression in its genome

To identify the closest relatives of the Saqqara leaf, we used Illumina sequencing reads available in the Sequence Read Archive to assemble the plastomes of 17 modern Asian and African date palms (including possible wild-origin individuals from Oman, Gros-Balthazard et al., 2017) and 17 individuals belonging to five closely related species (i.e. *P. atlantica*, *P. canariensis, P. reclinata*, *P. sylvestris,* and *P. theophrasti*; Table S2). To compare the outcome of our phylogenetic and population genomic analyses with results obtained by previous studies, our taxon sampling strategy is virtually identical to the one conducted by Flowers *et al.* (2019) and Gros-Balthazard *et al.* (2017). Maximum Likelihood (ML) phylogenetic analyses on full plastome alignments revealed that the Saqqara leaf is nested in a strongly supported clade (Likelihood Bootstrap Support [LBS]: 87) entirely composed of North African cultivated date palms and two accessions of *P. atlantica* (Fig. 1; Fig. S2), a disputed species currently restricted to Cape Verde (Gros-Balthazard et al., 2020a). The North African clade is itself nested in a clade (LBS 99) of *P. sylvestris* samples, a species found from Pakistan to Myanmar, long hypothesized to be the closest relative of *P. dactylifera* (Barrow 1998; Pintaud et al., 2013; Gros-Balthazard et al., 2020a). Asian *P. dactylifera* (LBS 100) form a clade that is sister to the above-described clade of African *P. dactylifera* plus *P. sylvestris* (LBS 100).

To determine the genomic affiliation of the nuclear genome of the Saqqara sample to either North African or Asian modern *P. dactylifera* populations, we conducted a model-free principal component (PCA) and a model-based clustering analysis using genotype likelihoods derived from the nuclear genomes of all accessions, the latter assuming two to eight ancestral populations. We conducted genomic clustering analyses using two genomes of *P. dactylifera* as reference with different levels of completeness and contiguity to account for read mapping and missing data biases (Skotte et al., 2013; Günter and Nettelblad, 2019; see *Methods*). Regardless of the reference genome used, with four populations the Saqqara genome grouped with populations of *P. dactylifera* with Asian ancestry. North African individuals of *P. dactylifera* and *P. atlantica* shared most of their genome with Asian modern date palm populations albeit with a relatively small proportion of their genome admixed with *P. theophrasti* (Fig. 2; Fig. S4), thus supporting previous findings (Flowers et al., 2019). When assuming five ancestral populations, *P. dactylifera* segregated into two populations with African and Asian ancestry, respectively (Fig. 2; Fig. S4). Here, the clustering indicated that the majority (90-98%) of the analysed nuclear sequences from the Saqqara genome displayed components of North African domesticated *P. dactylifera* individuals and Cape Verde’s *P. atlantica*, whilst the remaining 1-10% could be traced to both domesticated and wild Asian *P. dactylifera* individuals (Fig. 2; Fig. S4). Allele sharing between *P. dactylifera* and *P. sylvestris* was also evident in clustering analyses with four and five populations, also supporting previous findings by Flowers et al. (2019). The PCA revealed similar results to those obtained by the model-based clustering analyses. Regardless of the reference genome used, the covariance matrices inferred from ~27,000 to ~35,000 filtered sites placed the Saqqara date palm genome closest to modern North African date palm individuals in a cluster made of accessions of *P. dactylifera* (Fig. 2c,d).

**Figure 2.**
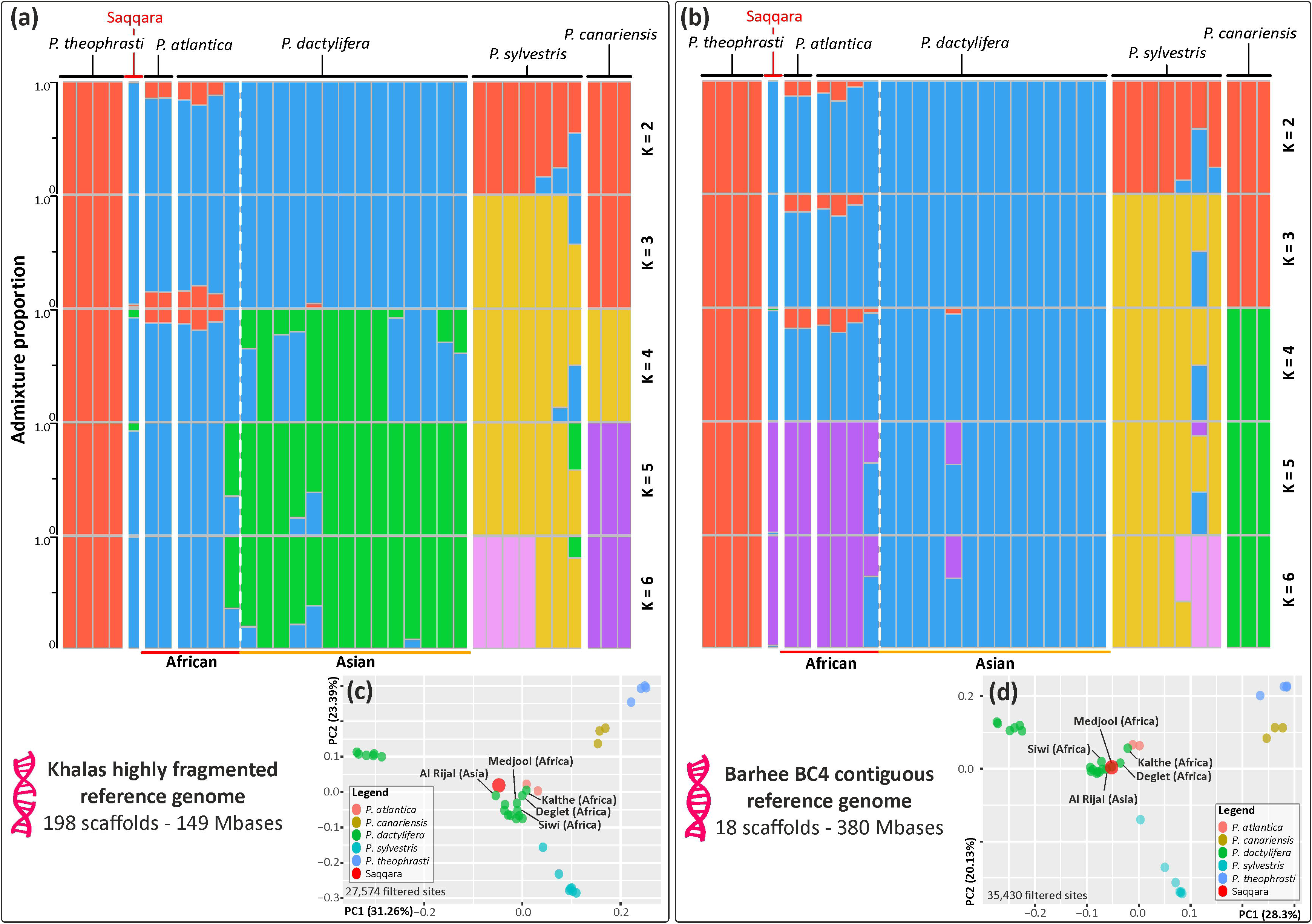
Genome ancestry of the Saqqara specimen. Population structure and principal component analyses (PCA) based on estimated nuclear GLs derived from a highly fragmented (**[a, c]**, GCA000413155.1) and a highly contiguous reference genome (**[b, d]**, GCA0009389715.1). Structure analyses with population number (K) from 2-6 (**a, b**) show admixture amongst wild and cultivated date palm populations, including the Saqqara leaf, and closely related *Phoenix* species. The geographical origin of modern individuals of *P. dactylifera* is provided at the bottom of the plot. Detailed cluster and delta likelihood values from K 1-8 are provided on Fig. S4. Covariance matrices derived from PCA in (**b, d**) reveal a close affinity of the Saqqara specimen with modern individuals of North African *P. dactylifera* and the Cape Verde’s *P. dactylifera*. The remaining individuals of *P. dactylifera* not labelled in the plots belong to Asian populations.

We conducted introgression tests (i.e. D-statistics) to trace gene exchange between the Saqqara leaf in relation to the modern date palms and the closely related *P. atlantica*, *P. sylvestris* and *P. theophrasti*, using *P. reclinata* as an outgroup based on previous studies (Flowers et al., 2019; Fig. 3; Fig. S3). To account for the differences in sequencing coverage obtained from modern individuals and the ancient Saqqara genome, these topological tests were conducted using two approaches tailored to separately evaluate individuals (i.e. by sampling one base from reads of one individual per population) and populations (i.e. by considering all reads from all individuals in each population; Soraggi et al., 2018). Both approaches were also implemented using two reference genomes to account for potential sequence biases (Günther and Nettelblad, 2019; see *Methods*). Analyses considering individuals separately and populations (regardless of the genome of reference) gave virtually identical results regarding the relatedness of the Saqqara leaf.

**Figure 3.**
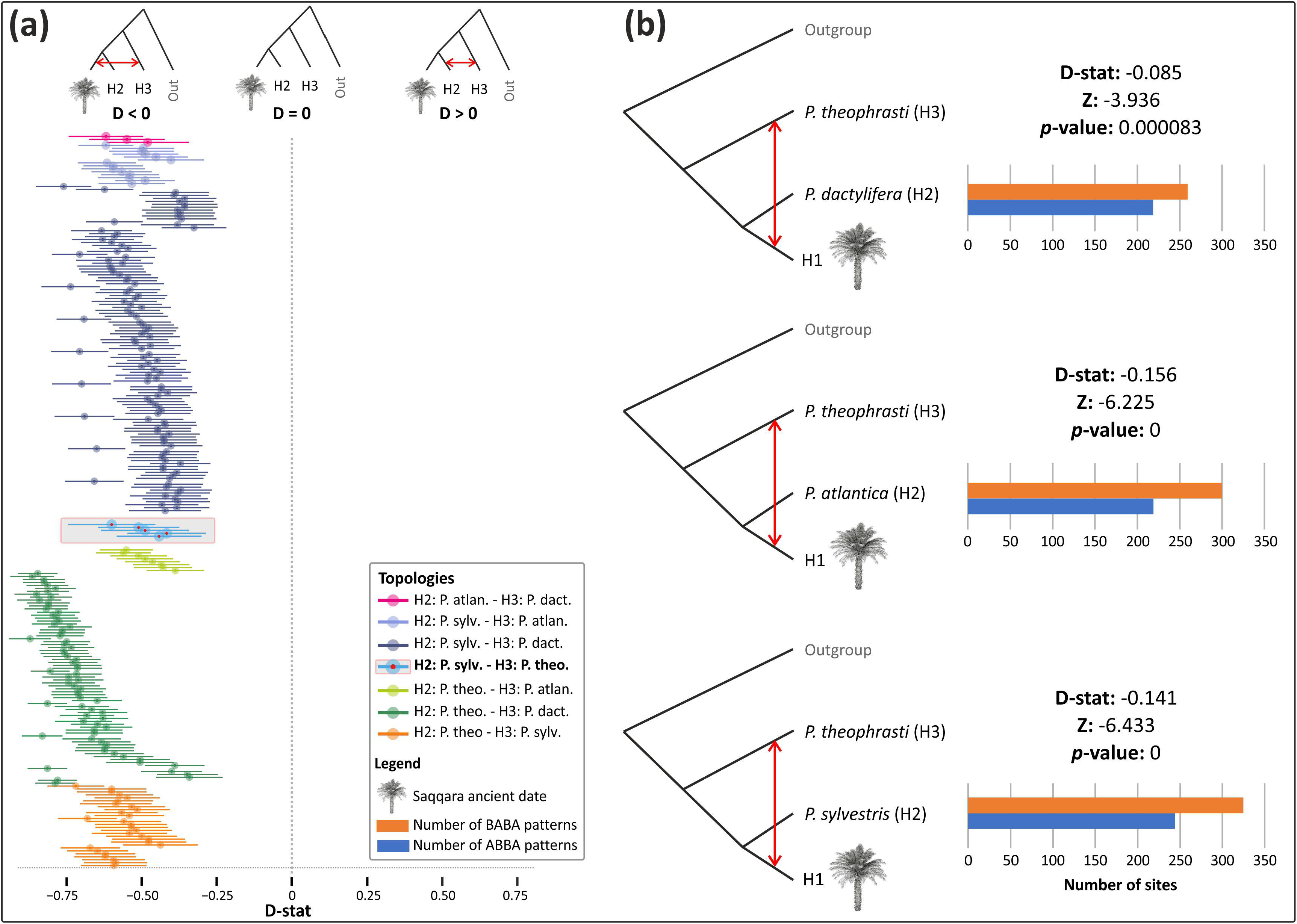
Introgression of the Saqqara leaf with modern individuals of *P. theophrasti* inferred from nuclear bases. (**a**) Results of D-statistic analyses derived from nuclear genotype likelihoods (GLs) for the Saqqara date leaf amongst date palm populations and closely related species (*P. atlantica*, *P. sylvestris* and *P. theophrasti*), with *P. reclinata* fixed as the outgroup, using as a reference a highly fragment reference genome. The outcome of all possible permutations conducted during the D-statistic test between all individuals sampled in this study are provided in Table S3 and Figure S3. Table S4 provides the outcome of D-statitstic conducted on all possible combinations between all individuals and using a contiguous reference genome. (**b**) Three instances of D-statistic analyses for the Saqqara date leaf conducted amongst populations of date palms and closely related species and a highly fragmented reference genome supporting gene flow between *P. theophrasti* and the Saqqara date leaf. The outcome of all possible permutations between populations and of analyses conducted using a contiguous reference genome is provided in Tables S5 and S6.

Introgression tests between modern individuals and the Saqqara leaf involved the evaluation of 39,280 and 503,386 nuclear bases, with an average of 81 and 586 bases per analysis using highly fragmented and contiguous genome assemblies as reference, respectively (Tabs. S3, S4). We found no signal of introgression from *P. sylvestris* in the Saqqara leaf nuclear genome as inferred from the analysis conducted on a highly fragmented genome. However, when computing D (Saqqara, *X*, *P. sylvestris*; *P. reclinata*, where *X* is *P. atlantica* or *P. dactylifera*) using as a reference the contiguous genome assembly, the Saqqara sample shared more derived alleles with *P. sylvestris* than with *X* (Table S4). Signal of introgression from *P. theophrasti* in the Saqqara sample was evident in analyses conducted on both highly fragmented and contiguous reference genomes. Here, when computing D (Saqqara, *P. sylvestris*; *P. theophrasti*, *P. reclinata*), the Saqqara sample shared more derived alleles with *P. theophrasti* than with *P. sylvestris* (Z < −3.1 and Z < −4.8 for highly fragmented and contiguous reference genome, respectively; Fig. 3A & Tabs. S3, S4).

Population tests using the highly fragmented genome of reference evaluated 3,472 to 9,469 bases, with an average of 192.93 and 527.6 bases per analysis considering polymorphic and non-polymorphic sites in the outgroup, respectively (Tabs. S5, S6). In contrast, analyses based on the continuous reference genome assessed 10,400 to 16,565 bases, with an average of 577.82 and 920.3 bases per analysis considering polymorphic and non-polymorphic sites in the outgroup, respectively (Tabs. S5, S6). Altogether, the population test analyses revealed results consistent with introgression analyses conducted at the individual level, regardless of the genome of references employed, thus providing support for the past occurrence of gene flow between the Saqqara leaf and *P. theophrasti*. Here, when computing D (Saqqara, *X*; *P. theophrasti*, *P. reclinata*, where *X* refers to either *P. sylvestris*, *P. atlantica*, or *P. dactylifera*), the Saqqara leaf genome shared more derived alleles with *P. theophrasti* than the latter with any of the other closely related taxa evaluated (Z < −3.93 < −6.9 & Z < −6.73 < −17.08 for highly fragmented and contiguous reference genomes, respectively; Fig. 3B; Tabs. S5, S6). As for individuals-based tests, only the D-statistic analysis conducted on the contiguous genome revealed gene flow between the Saqqara leaf and *P. sylvestris* (with D[Saqqara, *P. atlantica*; *P. sylvestris*, *P. reclinata*] = Z < −5.7; Table S5).

## Discussion

Generating genomic data from plant archaeological remains of known origin and unequivocal age provides a unique window into the timing and sequence of plant crop domestication and diffusion processes (Estrada et al., 2018; Gutaker et al., 2017; Swarts et al., 2017; Scott et al., 2019). Archaeological evidence thus far has not informed us about the occurrence and timing of genetic exchanges between date palms and their wild relatives, and whether their distribution overlapped in the past (Flowers et al., 2019). Our study is the first to address this gap by applying archaeogenomic approaches to shed light on date palm evolutionary history. Though low in overall proportion, we retrieve sufficient genetic information from the endogenous aDNA, highlighting the potential for further genomic analysis using additional archaeological remains, from other species, places and times.

Comparisons of our plastid and nuclear topologies and population structure analysis conducted on two reference genomes representing contrasting levels of contiguity and completeness provide robust evidence for the genomic affiliation of the Saqqara leaf with modern North African *P. dactylifera* populations, as well as the occurrence of ancient gene flow between *P. dactylifera, P. theophrasti,* and *P. sylvestris*. The clustering of *P. sylvestris* with selected North African date palm cultivars in plastid phylogenies has been previously reported by several studies (Pintaud et al. 2013; Chaluvadi et al., 2019; Flowers et al., 2019; Mohamoud et al., 2019), thus opening the question of whether gene flow or ancestral polymorphisms are responsible for such patterns (Flowers et al., 2019). *Phoenix dactylifera* and *P. sylvestris* overlap their distribution ranges in north western India and Pakistan, they are interfertile and known to produce fertile hybrids (Newton et al., 2013), suggesting that gene flow between both species is plausible.

Recently, extensive sequencing of over 200 organellar genomes of *P. dactylifera* revealed that date palm cultivars contain four haplotypes that are tightly linked to the geographical origin of the cultivar (Mohamoud et al., 2019), but the time of their diversification is largely unknown. In particular, one major haplotype (NA1, see Fig. 2 in Mohamoud et al., 2019) reported for North Africa is thought to be highly divergent from the remaining three haplotypes and is shared with *P. sylvestris* (Mohamoud et al., 2019). The trace of gene flow between the Saqqara leaf and *P. sylvestris* detected by our introgression analyses suggests that the recurrent clustering patterns of individuals from both species in plastid phylogenies could be derived from one or several chloroplast-capture processes mediated by hybridisation. In addition, by confidently placing the Saqqara leaf plastid genome in the NA1 haplotype, we set for the first time a minimum age for the origin of this plastid subpopulation to *c.* 2,100 yrs BP.

In the Nile valley, date stones are recovered from at least seven archaeological sites in Egypt and three in Sudan spanning the Middle Kingdom (2,500-1,650 BC) to 2^nd^ intermediate period, but they are only commonly present from the New Kingdom (1,570-1,070 BC) onwards (see reviews in Murray 2000; Zohary et al., 2015; and database in Flowers et al., 2019). More limited evidence from the Old Kingdom (2700-2100 BC) includes findings of two date stones, and occasional fragments of other plant parts, from Giza (Malleson and Miracle 2018). Textual evidence from the Old Kingdom also refers to imported dates (Tallet 2017). As such, some potential early finds may represent either imports or cultivation (Gros-Balthazard et al 2020). Examples of date stones recovered from earlier predynastic sites, notably from El Omari and Hierakonpolis, might be intrusive and lack reliable context (R. Friedman, pers. com. 1 March 2020) (Flowers et al., 2019). There are also occasional (potential) examples of date palm leaves and fibre from various sites, mostly funeral contexts, from around 3,800 B.C.E. onwards (Vartavan et al., 2000).

Flowers et al. (2019) suggested date cultivation in Egypt was established between the Middle to New Kingdom periods based on presence/absence of date stones recovered from archaeological sites within their database. Alternatively, Gros-Balthazard et al (2020) argue that cultivation and cultural importance is only clear from the New Kingdom onwards. The new importance of date culture during the New Kingdom is also reflected artistically, for instance in garden scenes within tomb wall-paintings (Parkinson 2008). Further west of the Nile valley, evidence for date cultivation at Zinkekra in Libya is clear from the early first millennium B.C.E, and also provides an early example of oasis agriculture in North Africa (Pelling 2013, Van der, Veen, & Westley, 2010). Additionally, a datestone dating to c.1400-1300 B.C.E. from the Wadi Tanzzuft, some 400 km further south-west, suggests earlier evidence for mid to later 2nd millennium date cultivation in the central Sahara (Mattingly and Wilson 2010, di Lernia and Manzi 2002). It may be that small scale cultivation was a precursor to a more widespread practice. This might reflect the time and expertise it takes to establish date palms effectively via clonal propagation rather than from seed, or to develop an adequate ratio of male/female trees and the practice of deliberate fertilisation (Bacon 1948). Thirdly, improved water management techniques were developed by the New Kingdom (Murray 2000).

Evidence demonstrating ancient date palm use and exploitation is recorded by reliable finds of date seeds around 5,500 B.C.E. from the Arabian Peninsula (Beech et al., 2003) and around 4,000 B.C.E. from Mesopotamia (Gillet, 1981; Tengberg, 2012). Date palms also occurred in Jordan from between 4,800-4,250 B.C.E. and in Israel from around 3,500-3,200 B.C.E. (Zohary et a., 2015). Taken together, these findings suggest an origin of date palm domestication around the Persian Gulf or Mesopotamia followed by the subsequent spread of the crop into North Africa. Several studies show two genetically distinct populations of domesticated *P. dactylifera* in North Africa and the Middle East (Gros-Balthazard et al., 2017; Gros-Balthazard et al., 2020b). The diversity of African date palms is, however, higher than expected following such a founder event, raising the possibility of an independent origin of date palm domestication in Africa and/or further introgression of genetic material from other *Phoenix* species (Hazzouri et al. 2015; Gros-Balthazard et al., 2017).

The study by Flowers et al. (2019) provided some explanation for the origin of the elevated genetic diversity in North African *P. dactylifera* by showing that parts of their genome were most closely related to *P. theophrasti*, a species currently restricted to the coastal areas of Crete, the Aegean islands and Turkey (Boydack, 2019). Following the only known archaeobotanical record of *P. theophrasti* found in northern Israel, the timing of such genetic exchange was hypothesised to have occurred ~7,500 yrs ago when the geographic ranges of the two species may have overlapped (Flowers et al., 2019). However, the authenticity of such an archaeological macrofossil is questionable because its identification relied solely on morphological comparisons of seed size, a character proven to be labile in wild and domesticated date palms (Gros-Balthazard et al., 2017; J. Dransfield, pers. com. 20 Feb 2020) and informative only when studied using statistical frameworks (Terral et al., 2012; Gros-Balthazard et al., 2017). Thus, before our study there was no concrete evidence supporting a minimum age for the genetic exchange between North African populations of *P. dactylifera* and *P. theophrasti.* Modern hybrid zones between *P. dactylifera* and *P. theophrasti* are known, and the species are known to hybridise in botanical gardens and plantations (Gros-Balthazard et al., 2013). The small but consistent proportion of derived alleles shared between the Saqqara date palm and *P. theophrasti* identified here (Figure 3) provides evidence that hybridisation between these species had already occurred before ~2,100 yrs BP. Nevertheless, caution is required when interpreting our D-statistics because the number of homologous bases analysed (ranging from hundreds to thousands; Tabs. S3-S6) represents only a small fraction of the nuclear genome. A better representation of the Saqqara nuclear genome, ideally attained through target capture to increase the proportion of endogenous nuclear DNA, would help to further define the proportion of the Saqqara genome that has been inherited from *P. theophrasti* or any other wild relative. In spite of the ancient timing of this introgression, the regions inherited from *P. theophrasti* comprise as much as 18% of the modern nuclear genome of North African domesticated date palm (Flowers et al., 2019). This high proportion suggests that such alleles may confer an advantage over their date palm homologs, enabling them to persist in the date palm genome despite the absence of a current contact zone between both species. Alternatively, the early implementation of clonal propagation of offshoots derived from hybrid individuals could have contributed to the survival of admixed genotypes. Archaeological evidence supporting such agricultural practice, however, is lacking.

## Conclusion

As the first documented instance of successful retrieval of ancient DNA from date palm, our study provides a timestamp for the occurrence of a key introgression process in the evolutionary history of this culturally and economically important crop. Our plastid and nuclear topological frameworks together with the genomic composition analysis involving different reference genomes, levels of contiguity and genomic representations, consistently indicate that the Saqqara palm belongs to the same clade as the modern North African *P. dactylifera*, and that by c. 2,000 yrs ago, this lineage had already undergone considerable genetic differentiation from Asian date palms, including from wild date palm populations. Our results also suggest a minimum date for ancient interspecific gene flow between North African *P. dactylifera, P. sylvestris* and *P. theophrasti*.

Our study highlights how an integrated approach comprising genomics, phylogenomics and archaeobotany can yield new insights on the timing and processes involved in the domestication and diffusion of one of the oldest fruit crops. The research showcases the importance of safeguarding and curating biological collections, which are now providing a multitude of uses in evolutionary research that could never have been dreamt of at the time samples were collected. Our results provide the temporal framework of date palm introgression with *P. theophrasti*, and a sound base for further biogeographical and molecular dating studies that could include a larger sampling of modern and ancient populations to clarify a) whether gene flow with other wild relatives also occurred and b) where and in what environmental conditions these processes took place.

## Supplementary Information

Supplemental Information includes four figures and six tables.

## Author Contributions

O.A.P.E., S.B., M.N., M.G.-B. and W.B. conceived the study. M.G.-B., J.F., O.A.P.E., S.B., M.H., M.Pu., A.S.T.P., and R.G., designed in-silico analyses. M.N., P.R., and M.L. conducted archaeobotanical research. P.P., B.G., M.H., M.Pr. and I.H conducted lab work. O.A.P.E., T.W., R.S., R.D., and D.B. conducted in-silico analyses. O.A.P.E., M.N., P.R., S.B., S.D., I.L., M.Pr., S.S.R. G.C., and W.R. wrote the manuscript, with contributions from all authors.

## Acknowledgements

We would like to thank Valentina Gasperini (The British Museum) for her advice on the Saqqara excavation contexts, and Renée Friedman (Director of excavations at the Predynastic site of Hierakonpolis) for her advice on the reliability of date stone archaeological findings and the need for re-evaluation of date leaf finds.

