## Supplementary figures and images for "Archaeogenomics of a ~2,100-year-old Egyptian leaf provides a new timestamp on date palm domestication"

### Fig S1

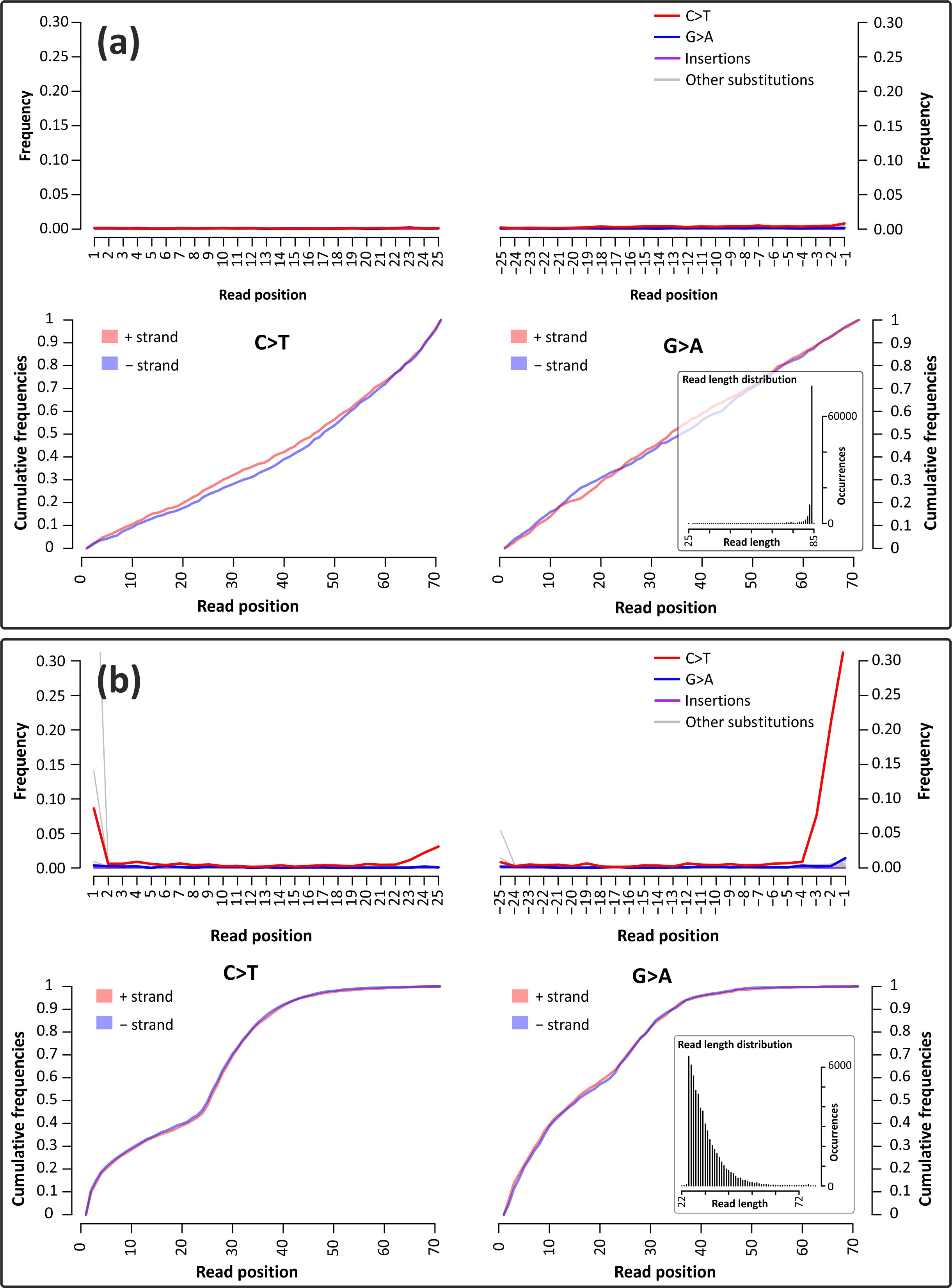

### Fig S2

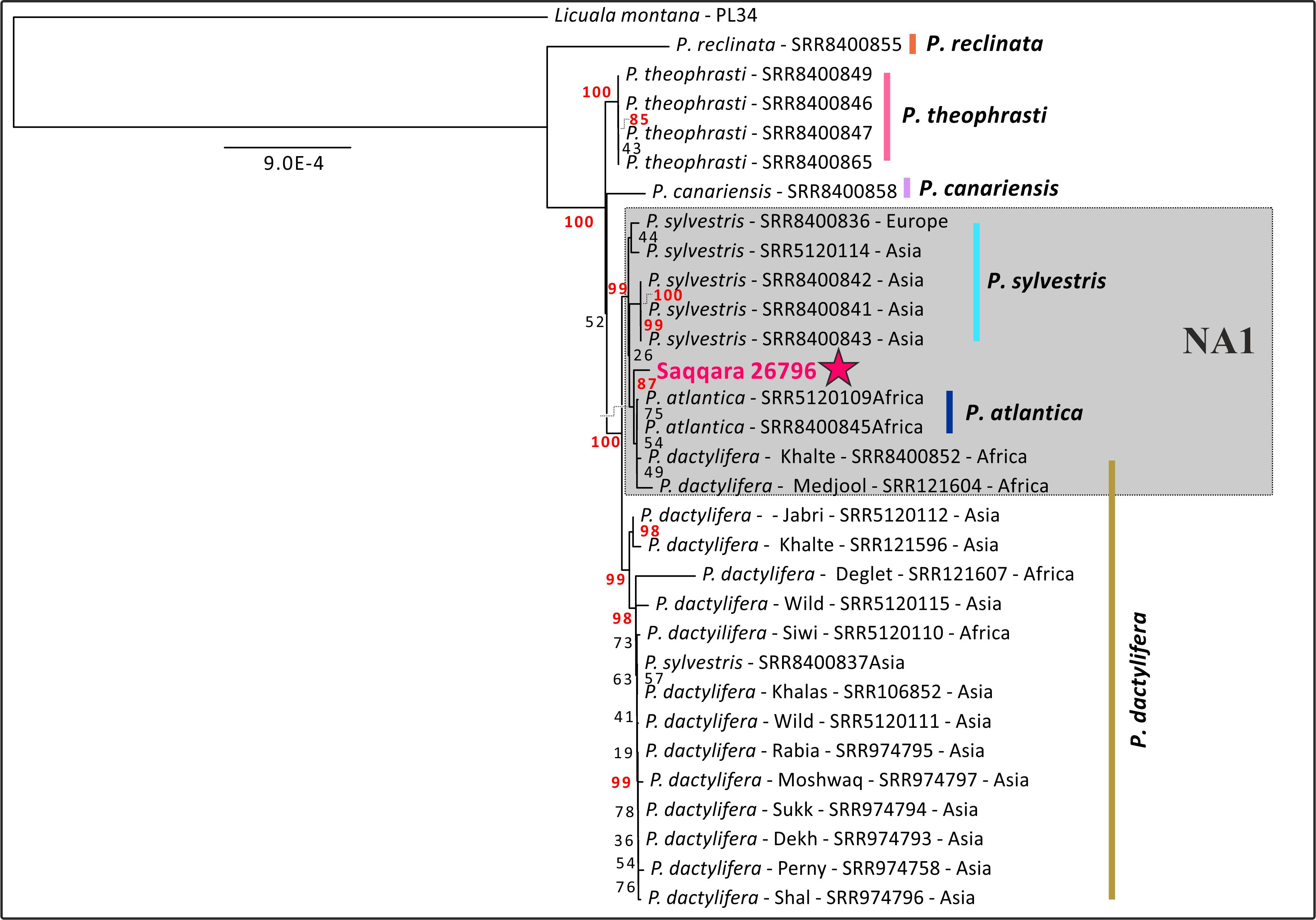

### Fig S3

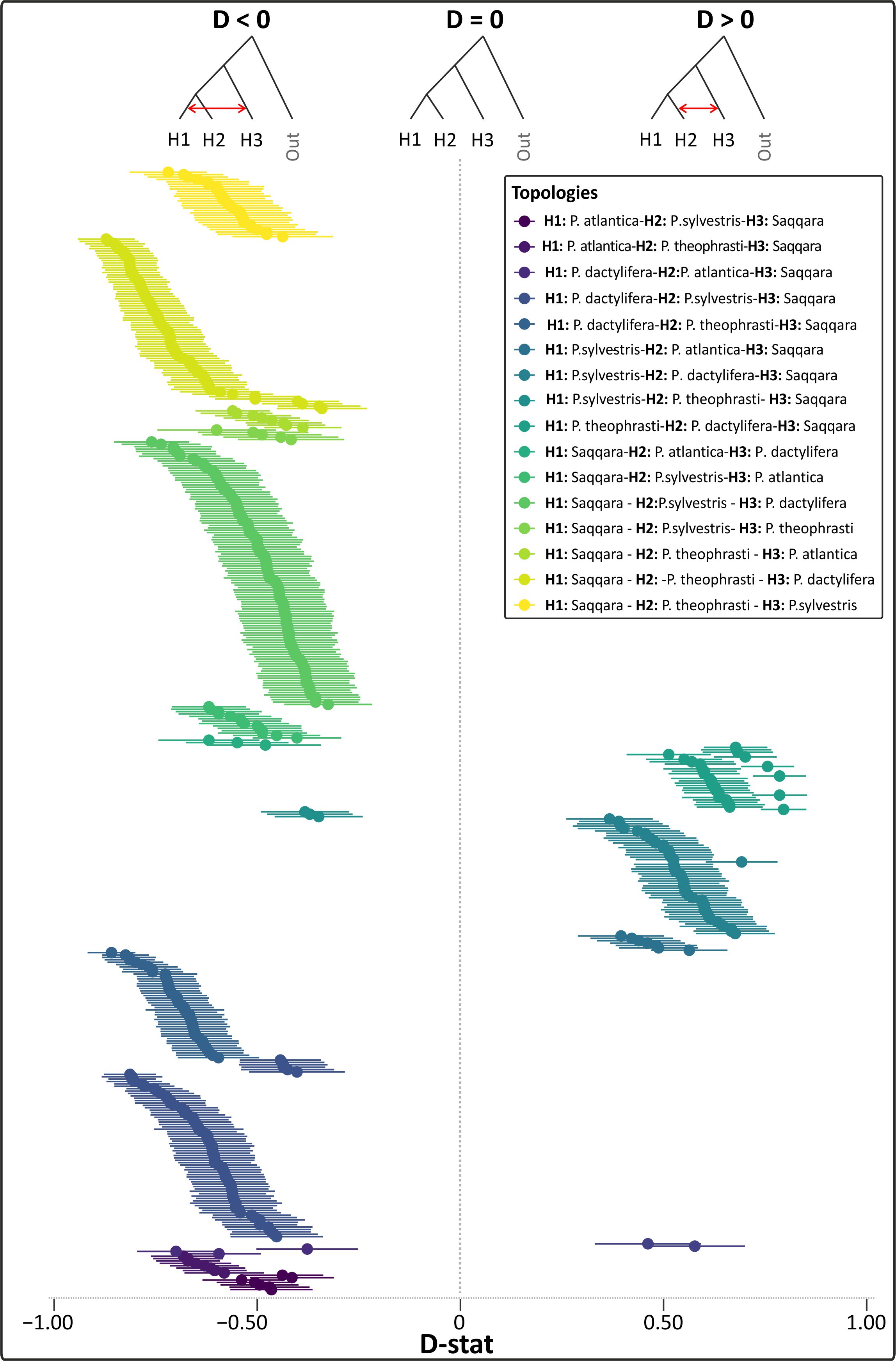

### Fig S4

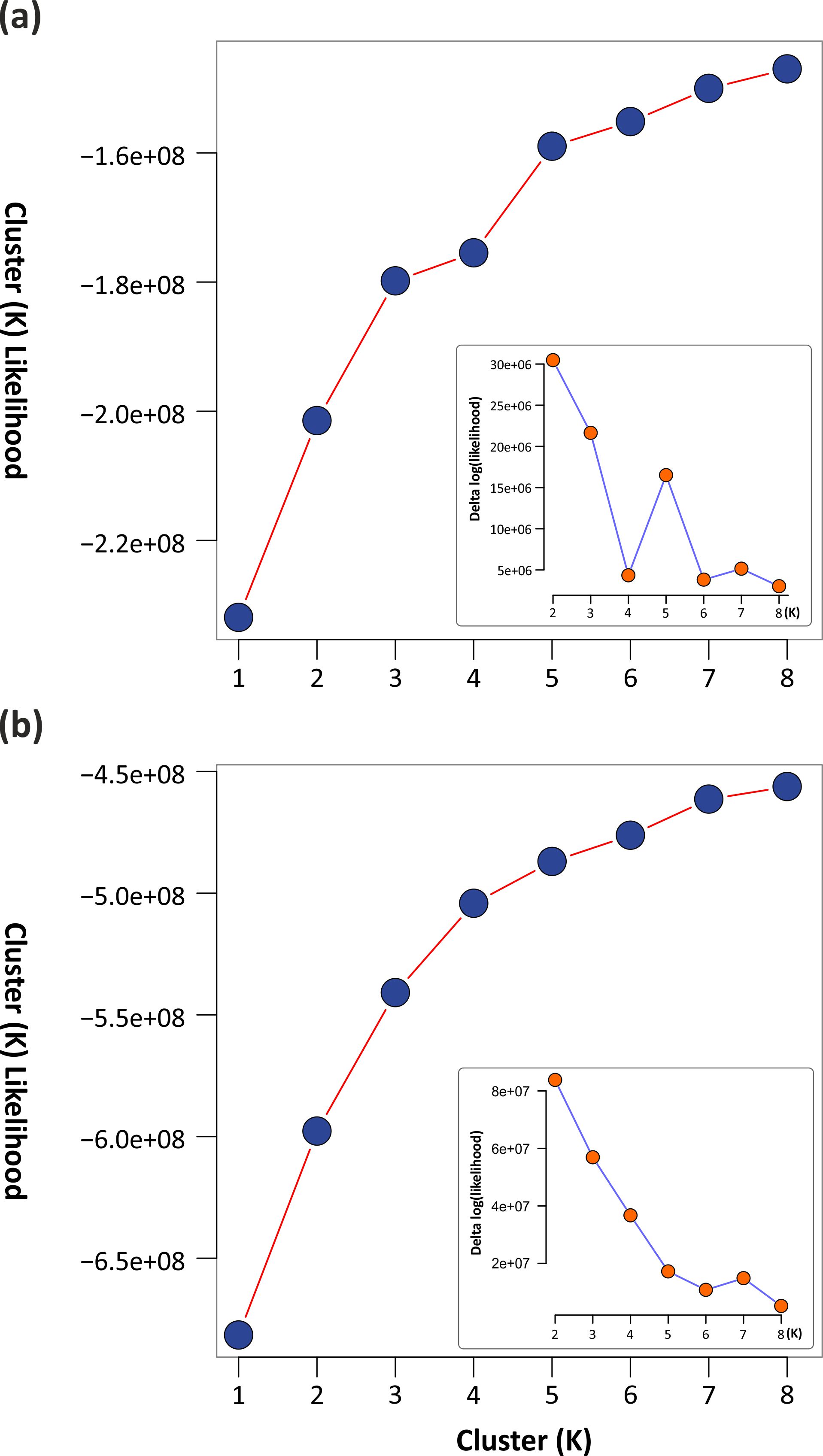
